# Hawaiian Fish Sounds and their Potential as Acoustic Ecological Indicators on Coral Reefs

**DOI:** 10.64898/2026.08.10.744083

**Authors:** Erika Berlik, Marc S. Dantzker, Symeon Delikaris-Manias, Matthew T. Duggan, Aaron N. Rice

## Abstract

Coral reef monitoring needs scalable, non-invasive tools to complement resource-intensive traditional survey methods. Passive Acoustic Monitoring (PAM) offers a promising supplement, but its effectiveness is limited by the difficulty of attributing recorded sounds to species outside of previously well-characterized taxa. Using Omnidirectional Underwater Passive Acoustic Cameras (UPAC-360), we identified sounds from 31 reef fish species across 14 families on the Kona coast of Hawai’i Island, including 13 not previously documented as soniferous. By releasing video and audio specimens, we have created the largest open-access collection of in-situ reef fish sounds to date for the Pacific. A subset of acoustically distinctive taxa—such as Hawaiian Dascyllus (*Dascyllus albisella*), Lei Triggerfish (*Sufflamen bursa*), soldierfishes (*Myripristis spp.*), wrasses, and herbivorous grazers—were identifiable in PAM recordings through manual acoustic and spectrogram review. Through identifying particular sounds linked to species with different ecological roles, these sounds have the potential to serve as indicators of reef function to increase the information and value coming from PAM surveys of Hawaiian and Pacific coral reefs.

## Introduction

Monitoring coral reef fish populations is fundamental to understanding ecosystem resilience and effectively managing fisheries and conservation areas (e.g., Carrillo-García and Kolb 2022; Edmunds 2024; Flower et al. 2017; MacNeil et al. 2015; Wolfe et al. 2020). There is an increasing use of diverse technologies to augment conventional monitoring methods for evaluating coral reef communities (Apprill et al. 2023; Madin et al. 2019). As coral reefs decline globally (Hughes et al. 2017), the development of noninvasive and generalizable monitoring tools is critical for guiding conservation and management interventions (Edmunds 2024).

Passive Acoustic Monitoring (PAM) has addressed a wide range of reef management questions, including biodiversity surveys (Lin et al. 2021; Mahale et al. 2023; Minier et al. 2025; Ozanich et al. 2021), fisheries assessments (Rountree et al. 2006), habitat integrity (Freeman and Freeman 2016; Jarriel et al. 2024), MPA efficacy (Haver et al. 2018; Ruiz 2026), and anthropogenic impacts (Au et al. 2012; Duane et al. 2023; Ferrier-Pagès et al. 2021; Kaplan et al. 2018). Despite the recognition of the ubiquity, abundance, and importance of fish sounds on coral reefs (Hodson et al. 2025; Lobel et al. 2010; Looby et al. 2023; Parmentier et al. 2021; Rice et al. 2022) and the increasing understanding that most fish species likely make and use sound for communication (Dantzker et al. 2025a; Looby et al. 2023; Rice et al. 2022), the lack of species-specific validated sounds constrains the analytical insight and impact of the application of passive acoustics (for example, see Figure 1 in Dantzker et al. 2025a). When sounds cannot be attributed to a species or higher taxon, only limited measures can be extracted, such as noise levels (Duane et al. 2023; Haver et al. 2018; Kaplan et al. 2018; Kaplan et al. 2015; McKenna et al. 2021), acoustic indices (Bertucci et al. 2016; Harris et al. 2016; Minier et al. 2025; Williams et al. 2022), or unidentified fish sound classes (Duane et al. 2026; Jarriel et al. 2024; Lin et al. 2021; Williams et al. 2025). Noise levels are relevant measures for measuring anthropogenic noise; however, other abstract or computational measures of biological sounds often do not meet the specificity needs that managers and regulators require for making species- or habitat-level decisions (Flower et al. 2017; Sugai et al. 2026). The emerging ability to connect sounds to identified fish species (Dantzker et al. 2025a; Dantzker et al. 2025b; Mouy et al. 2023) provides the opportunity to significantly advance coral reef PAM for providing actionable biodiversity, conservation, and management information to stakeholders. Even short-duration PAM recordings or sounds from individual identified fish species from coral reefs can be valuable for applying acoustics-based reef fish monitoring and management in places of need (Ibrahim et al. 2024; Pyć et al. 2021; Tricas and Boyle 2021); this is particularly true in regions like the Hawaiian Islands where acoustic monitoring infrastructure already exists for other purposes (Fig. 1) (Au et al. 2012; Duane et al. 2023; Duane et al. 2024; Freeman and Freeman 2016; Haver et al. 2018; Havlik et al. 2022; Kaplan et al. 2018; Lammers et al. 2023; McKenna et al. 2021; McKenna et al. 2024).

**Figure 1.**
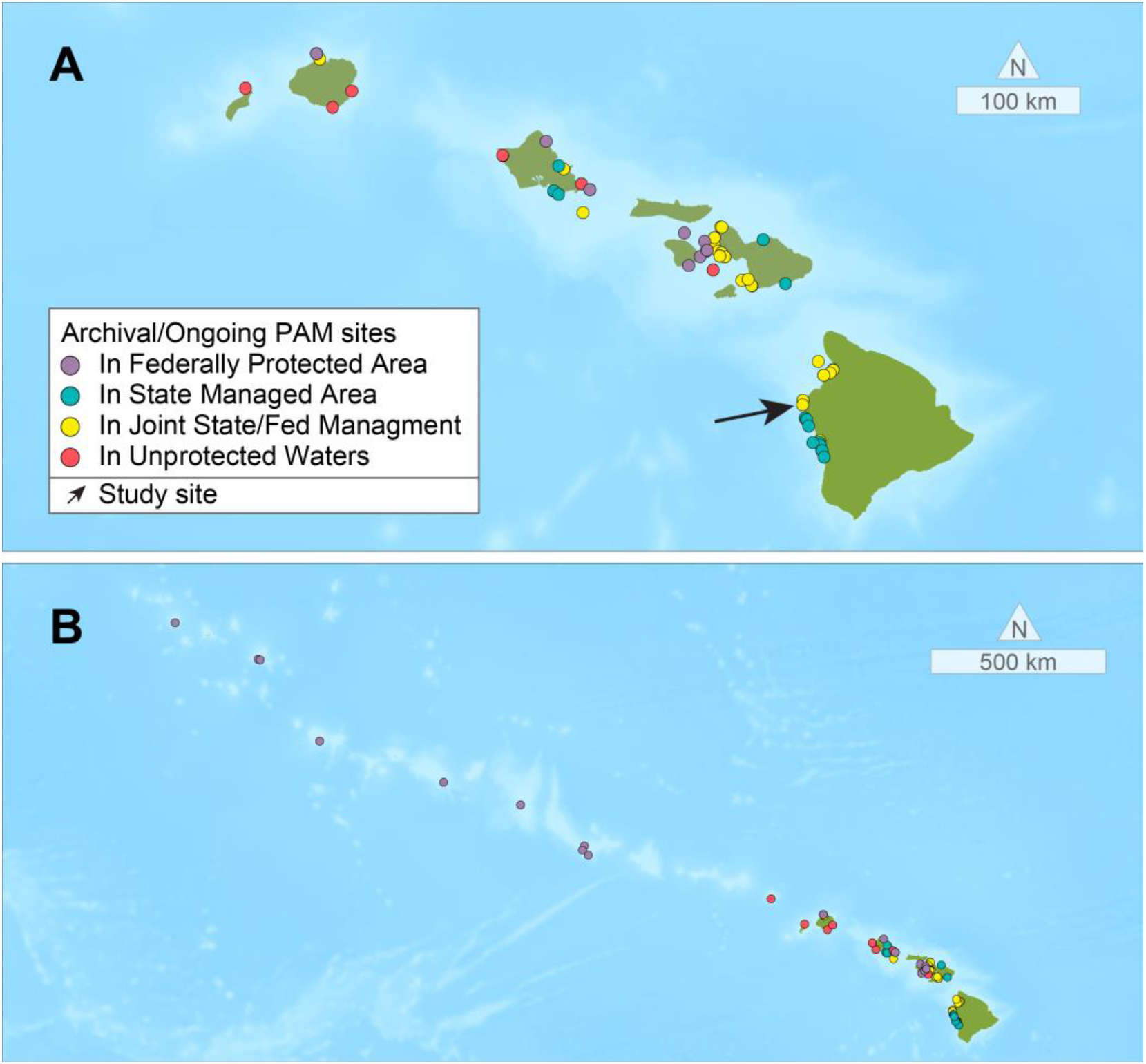
Map of existing PAM recording sites on the Hawaiian archipelago overlapping with shallow and mesophotic coral reefs. Each point on the map represents a known site at which PAM recordings exist on the main Hawaiian Islands (A) and the remainder of the Hawaiian archipelago (B). The location for our recording effort is marked with an arrow (A). Not all recordings will be comparable, as they vary in deployment methods, habitats, depths, and duration (ranging from days to years) among other factors. Sensor locations are from previously publications (Duane et al. 2024; Freeman and Freeman 2016; Haver et al. 2018; Havlik et al. 2022; Heenehan et al. 2017; Kügler et al. 2024; Lammers et al. 2023; Madrigal et al. 2024; McElligott and Lammers 2021; Thode et al. 2021) and NOAA’s NCEI Passive Acoustic Data Portal (https://www.ncei.noaa.gov/maps/passive-acoustic-data/).

Coral reefs in Hawai’i are among the most intensively monitored reefs, representing an essential natural and cultural resource (Costa and Kendall 2016; Friedlander et al. 2008; Jokiel 2008). Hawai’i has a significant amount of reef area that is managed by state and/or US federal agencies (e.g. National Oceanic and Atmospheric Administration, Department of Defense, Department of the Interior), which often entail long-term monitoring programs either in the context of fisheries, habitat, or ecosystem-based fisheries management (e.g., Boehlert 1993; Costa and Kendall 2016; Greenfield 2003; Jokiel 2008; Smith 1993). Like coral reefs around the world, Hawaiian reefs are vulnerable to climate change and anthropogenic impacts (Andrello et al. 2022; Friedlander et al. 2008; Selkoe et al. 2008), and connecting PAM to ecosystem regime shifts through ecologically important species and their sounds may provide actionable conservation and management information..

PAM is already being used in marine monitoring initiatives across the Hawaiian archipelago and continues to be collected on the main and Northwestern Hawaiian Islands (Fig. 1). These studies have primarily focused on marine mammal species and noise level characterization (Au et al. 2000; Haver et al. 2018; Kaplan et al. 2018; Lammers et al. 2023; Parnell et al. 2024; Zurk et al. 2014). While several have included focal or unidentified fish sounds as intentional targets for recording (Freeman and Freeman 2016; Freeman et al. 2014; Lobel and Mann 1995; Mann and Lobel 1995; Mann and Lobel 1997; Mann and Lobel 1998; Ozanich et al. 2021; Tricas and Boyle 2014; Tricas and Boyle 2021), an improved understanding of the Hawai’i’s vocal fish species enables these recordings to become a powerful archival record from which we can draw new insights with new acoustic information (Duane et al. 2023; Duane et al. 2026; Freeman and Freeman 2016; Havlik et al. 2022; Kaplan et al. 2018; Lammers et al. 2023). Because of the extensive amount of protected or managed waters in the Hawaiian Archipelago, the majority of existing PAM study sites fall within areas under state and/or federal protection with active natural resource management. Back-labeling of acoustic data could reveal patterns in acoustic activity of protected and/or essential commercial or subsistence fisheries species (Duane et al. 2026).

In their foundational study, Tricas and Boyle (2014) published spectral and temporal measurements from recordings of 45 species made on closed-circuit rebreather using a camera fitted with a hydrophone. A subset of these (21) was also presented as spectrograms and waveforms. These descriptions lay essential groundwork for our understanding of soniferous fish species in the Pacific; however, neither a table of acoustic measures nor spectrograms are typically sufficient to allow accurate matching or cross-referencing to other PAM recordings.

In this study, we identify and make available sounds from 31 reef fish species. We estimate detection potential and “acoustic identifiability” of their calls and evaluate their roles as acoustic indicator species for monitoring fish assemblages and informing reef management through PAM. Specimens from these species have been published in an open-access library (https://www.fisheyecollaborative.org/library; Dantzker et al. 2025c), and are cross-referenced with other data portals such as FishSounds.net (Looby et al. 2026). The identifications (IDs) found here, alongside those previously released for the Caribbean, are now part of the largest open-access library of in-situ recordings of both Atlantic and Pacific reef-associated fishes.

With this short duration case study, we identify a conceptual and technical framework of how fish-focused PAM can scale to long term PAM datasets for examining patterns of biodiversity, changes in community composition, response to anthropogenic impacts, or MPA efficacy in Hawai’i and the broader Pacific.

## Methods

### General Approach to Sound Identification

We have described our technical approach (based on Delikaris-Manias et al. 2018) to ascribing sounds to fish species in detail elsewhere (Dantzker et al. 2025a). In summary, we deploy compact acoustic and video arrays to autonomously record coral reef ecosystems. Divers leave the area for extended periods meaning observer effects are minimized, and fish are more likely to exhibit a broader range of natural behaviors. We have described the devices as Underwater Passive Acoustic Cameras (UPAC-360), and they consist of a compact tetrahedral hydrophone array positioned around a 360° camera. The UPAC-360 records audio and video separately but the signals are synchronized using conspicuous external acoustic events (e.g. hydrophone taps, diver sounds, loud impulsive environmental sounds) that are audible on the video camera’s internal microphones and the hydrophone recordings. In post-processing, the video is converted into an equirectangular projection of the 360° visual field using Insta 360 Studio software. The acoustic array recordings are converted to first-order ambisonic signals. Using a combination of beamforming algorithms (CroPaC and MUSIC; Delikaris-Manias and Pulkki 2013; Schmidt 1986), we produce a visualization of the acoustic energy in the soundfield from the same orientation as the camera. When an individual fish, or a small group of the same species, is the only potential source within the peak of the Direction of Arrival (DOA), we can ascribe the sound to that species.

### Fieldwork

Between April 25 and April 27 of 2022, we deployed two UPAC-360 systems at coral reef sites along the Kona coast of Hawai’i. Over the three days, we deployed systems at six locations, between 20 and 23 meters deep, along a 20-meter section of reef slope at the base of the reef wall (19.737° N, 156.054° W) in Makako Bay near the Ellison Onizuka Kona International Airport (Fig. 1). This location is a popular dive site known as “Garden Eels.” It lies within the West Hawai’i Regional Fishery Management Area and the southernmost part of the Hawaiian Islands Humpback Whale National Marine Sanctuary. The deployment locations were near the bottom of a coral-covered volcanic ridge, near the interface with a broad sand patch. We chose this site because we found a concentration of active *Dascyllus albisella* (Hawaiian Dascyllus or the Hawaiian Domino Damselfish) nests with its conspicuous and well-described acoustic behaviors (Laboury et al. 2025; Lobel and Mann 1995; Mann and Lobel 1995; Mann and Lobel 1997; Mann and Lobel 1998) that we could target (Fig. 2). Over three days, on the two devices, we made 21 hours of recordings.

**Figure 2.**
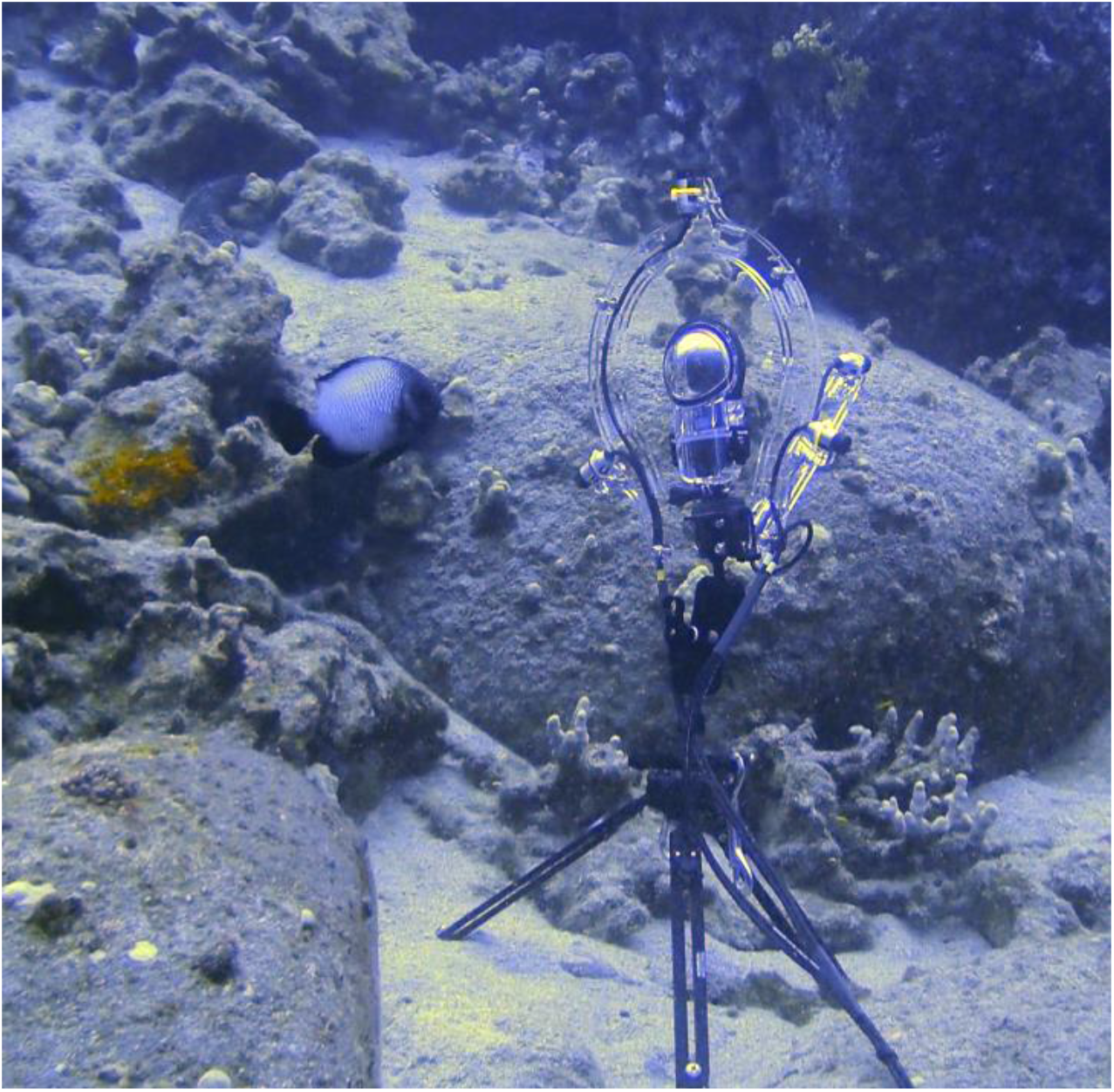
UPAC-360 sensor system deployed off the Kona Coast with adjacent displaying *D. albisella. Photo by ANR*.

### Hardware Details

For this study, the UPAC-360 was equipped with an Insta-360 X2 camera (Shenzhen, Guangdong, China) with a 5760×2280 resolution. The acoustic system is identical to Dantzker et al. (2025a), with prototype Aquarian Audio A5 hydrophones (Scientific, Anacortes, WA, USA), Ambisonic A format (at 48 kHz sample rate with 32 bit depth) recorded to a Zoom F6 (Zoom Corporation, Tokyo, Japan) in a Blue Robotics Inc. (Torrance, CA, USA) housing.

### Analysis

Analysis methodology largely followed the details outlined in Dantzker et al. (2025a) but with some workflow improvements. A continuous map of the acoustic energy’s spatial distribution was visualized in 8.33 msec increments with a powermap produced using C++ (version) of the SPARTA Toolbox (McCormack et al. 2018). The powermap visualization creates a 120fps video which is layered over the corresponding 30 fps video, along with a scrolling spectrogram. We used the Karta VR plugin V.5.7 (Kartaverse, https://kartaverse.github.io/) to reframe the 360° video and focus on individual sound events (Fig. 3).

**Figure 3.**
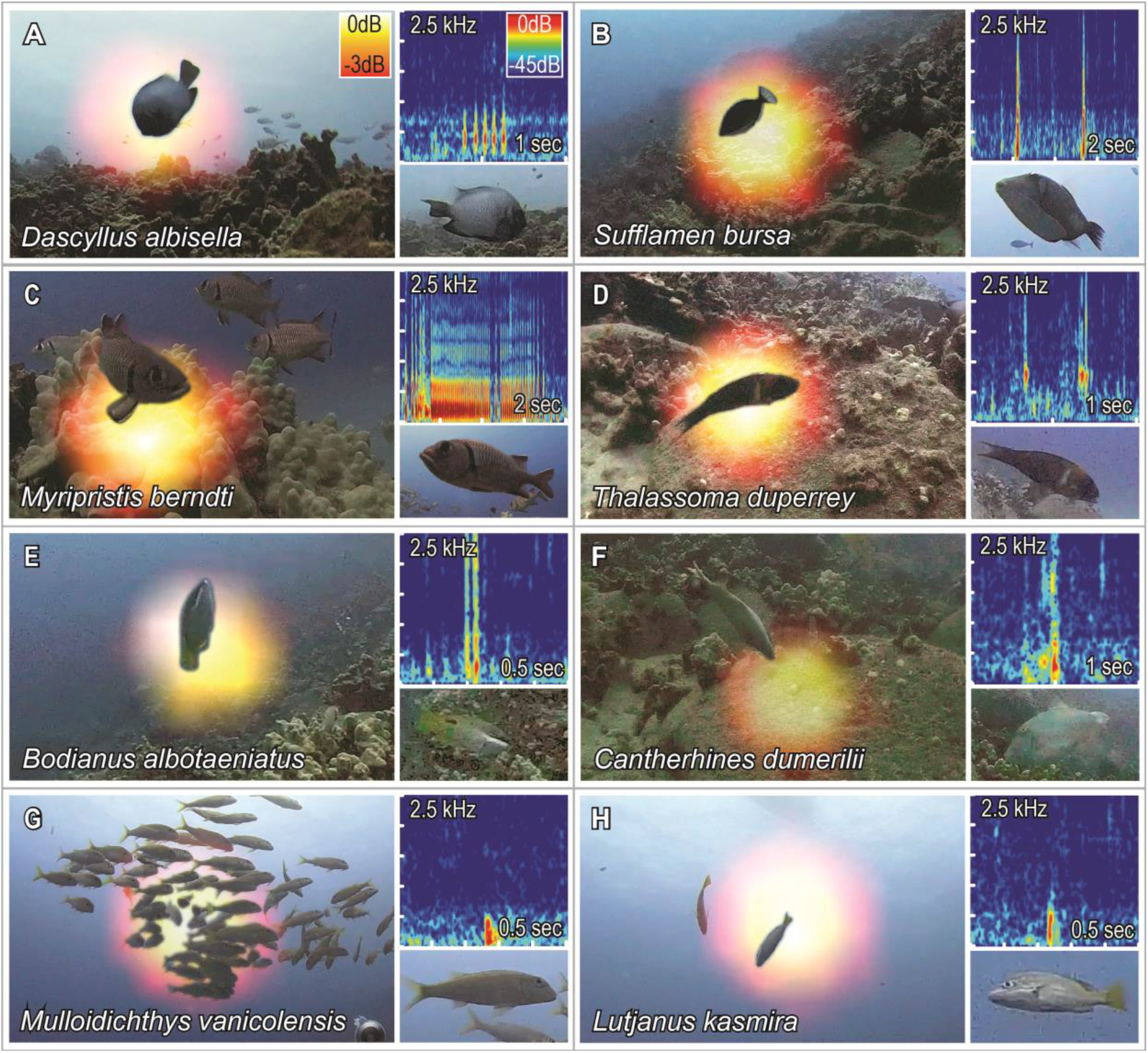
The sounds from Hawaiian reef fishes shown with the spectrogram and spatial audio powermap over a frame from the video sequence for: (A) *Dascyllus albisella*, Hawaiian Dascyllus, (B) *Sufflamen bursa*, Lei Triggerfish, (C) *Myripristis berndti*, Bigscale Soldierfish, (D) *Thalassoma duperrey*, Saddle Wrasse, (E) *Bodianus albotaeniatus*, Hawaiian Hogfish, (F) *Cantherhines dumerilii*, Whitespotted Filefish, (G) *Mulloidichthys vanicolensis*, Yellowfin Goatfish, and (H) *Lutjanus kasmira*, Bluestripe Snapper. The powermap layer has been removed where it overlaps fish to allow for a clear view of the sound-producing individuals.

The system can optimally localize sounds between 200-2500 Hz due the geometry of the array, and this is the frequency band that includes most known fish sounds (Lindseth and Lobel 2018; Rountree et al. 2006). The visual range of the UPAC-360 is limited by light, visibility, camera resolution, and the topography of the reef. It is not always possible to assign detected sounds to species. Visually cryptic and nocturnal species may produce sounds from within structures where they cannot be identified. In cases where the powermap overlaps multiple individuals of two or more species, the source of the sound can often be ascribed to one of those species using conspicuous behavioral cues such as courtship dances or agonistic chases that are known to be associated with sound production (Davidson et al. 2024; Oliver and Lobel 2013). However, with any ambiguity, we assign sounds to species only when the observation is corroborated by multiple other instances. Water temperature and salinity used for speed of sound calculation in spatial audio analyses were obtained from the PacIOOS data portal. (https://www.pacioos.hawaii.edu/water/model-salinity-hawaii/).

### Acoustic Identifiability

Each species was qualitatively scored on a scale of 1-3 for the identifiability of their sounds through aural and spectrographic analysis. This assessment was conducted by a single individual (EB) who had identified sounds from all previous UPAC-360 recordings (Dantzker et al. 2025a; Dantzker et al. 2025b). Species scored at 1 either produced sounds that were relatively subtle and indistinct from one another, or insufficient examples were collected to suggest their sounds may be identifiable in acoustic data. Species scored at 2 produced sounds that could be identified from audio but only to broad categories such as families or functional groups. Species scored at 3 produced frequent and unique vocalizations with consistent characteristics, often paired with unique behaviors.

### Discriminant Function Analysis

To corroborate the qualitative human visual analysis and classification of identifiability (described above) and enable quantitative replicability, we evaluated representative supervised features to identify if there is sufficient evidence to discriminate sounds to classes. Validated sounds were human-annotated with a close-fitting annotation to the signal (with minimal padding in time or frequency), and several acoustic features were measured in Raven Pro 1.6.5 (KLY-CCB 2022): Bandwidth 50%, Bandwidth 90%, Aggregate Entropy, Average Entropy, Maximum Entropy, Center Time, Center Frequency, Duration, 90% Duration, Peak Frequency, Peak Time, and number of pulses. These measurements were chosen to quantitatively and consistently characterize signals with minimal influence of annotation variability. For pulsed sounds, the number of pulses in each signal was measured manually from the waveform after applying a 50-2000 Hz bandpass filter in Raven. Human experts can readily distinguish fish communication sounds from feeding-related scraping sounds (such as from parrotfish or filefish) (Sartori and Bright 1973; Tricas and Boyle 2021), but we analyzed Maximum Entropy from feeding and communication sounds with a Kruskal-Wallis test to statistically corroborate these previous results. Maximum Entropy was used as a representative feature indicative of high-level differences, but other feature differences between classes were not explored further.

To inform the species-, genus-, or family-level classification of communication sounds, we conducted a Discriminant Function Analysis (DFA) in JMP Student Edition 19.1.3 (JMP Statistical Discovery LLC, Cary, NC), with a quadratic fitting to account for different within-group covariance matrices, using the Raven measurements as inputs for each sound. Since feeding sounds can be identified relatively easily, we focused our species-level statistical classification on only active communication sounds (vocalizations) in the DFA analysis. The DFA was used to test whether there is sufficient information and what parameters help to classify them into different taxa or functional groups. The sounds used for this analysis included only those contained within our library specimens. The specimens are intended to provide our best representatives across the acoustic repertoires of each species.

### Application in PAM Recordings

To demonstrate the application of the validated sound library in acoustic monitoring, we selected 30 minutes of audio collected by a single hydrophone from the UPAC-360’s array for manual spectrographic analysis. This section was taken from a day when the audio array was recording, but the video was not; thus, it is not included in the set of recordings used for identification. While it is not specifically from a standard PAM recorder, each single hydrophone recording is functionally akin to a PAM device. This recording was made on April 24, 2022 at a depth of 22 m at the study site. No video validation was used for this portion of the analysis to compare with video verified examples (Table 1). The spectrogram was manually annotated using Raven, and sounds were identified with the highest possible taxonomic resolution. Unidentifiable sounds were classified as unknowns.

**Table 1.**
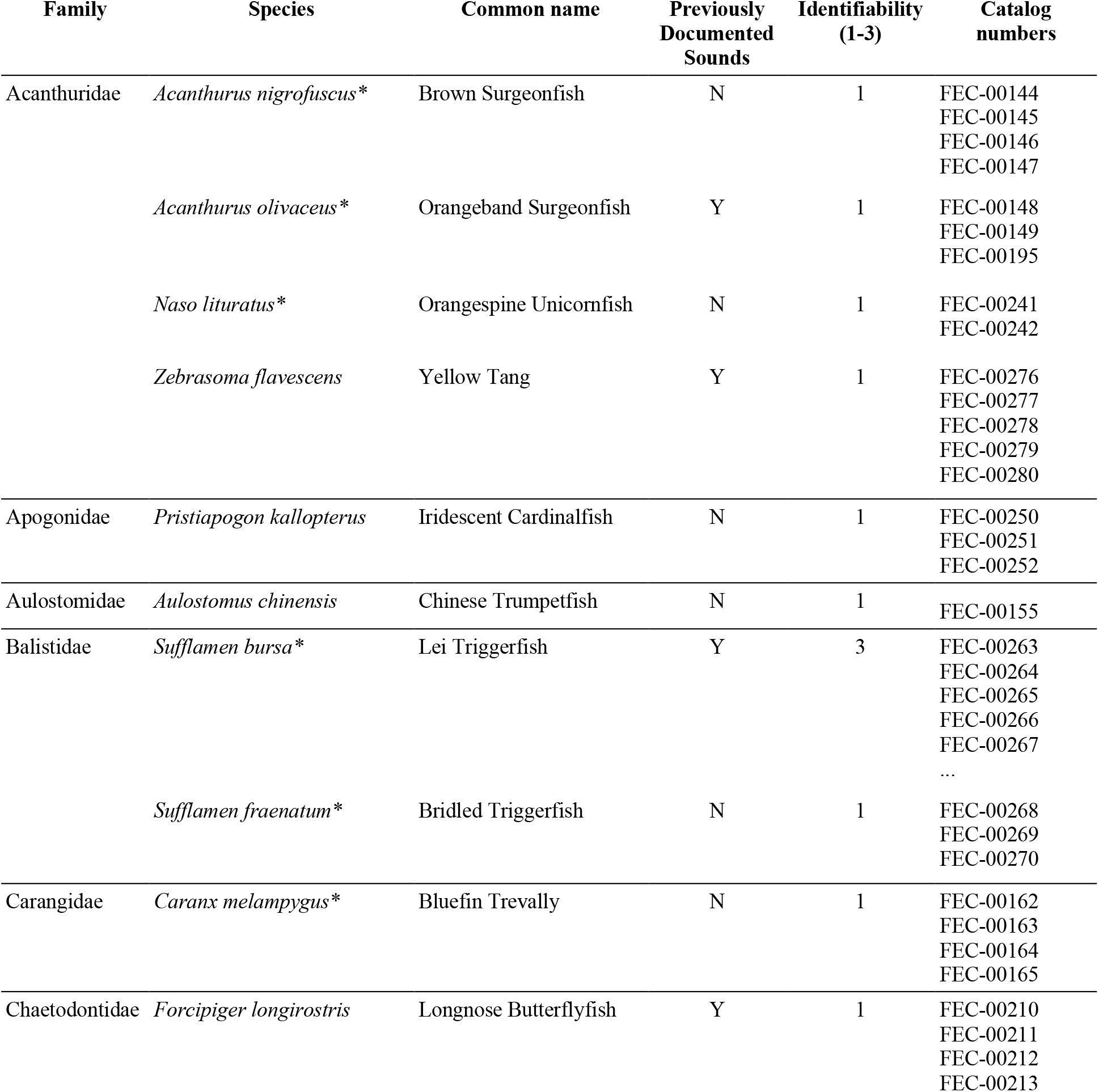
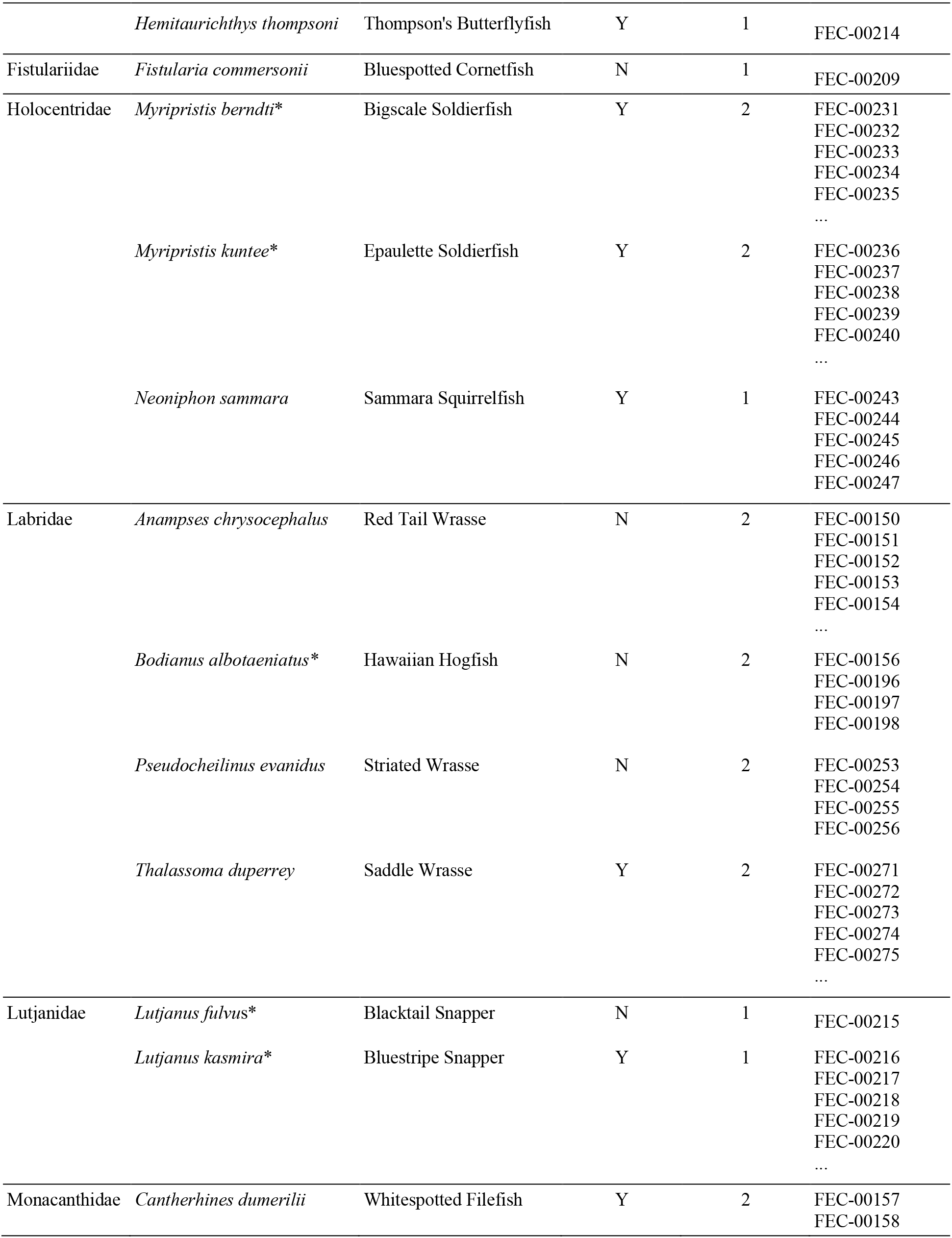
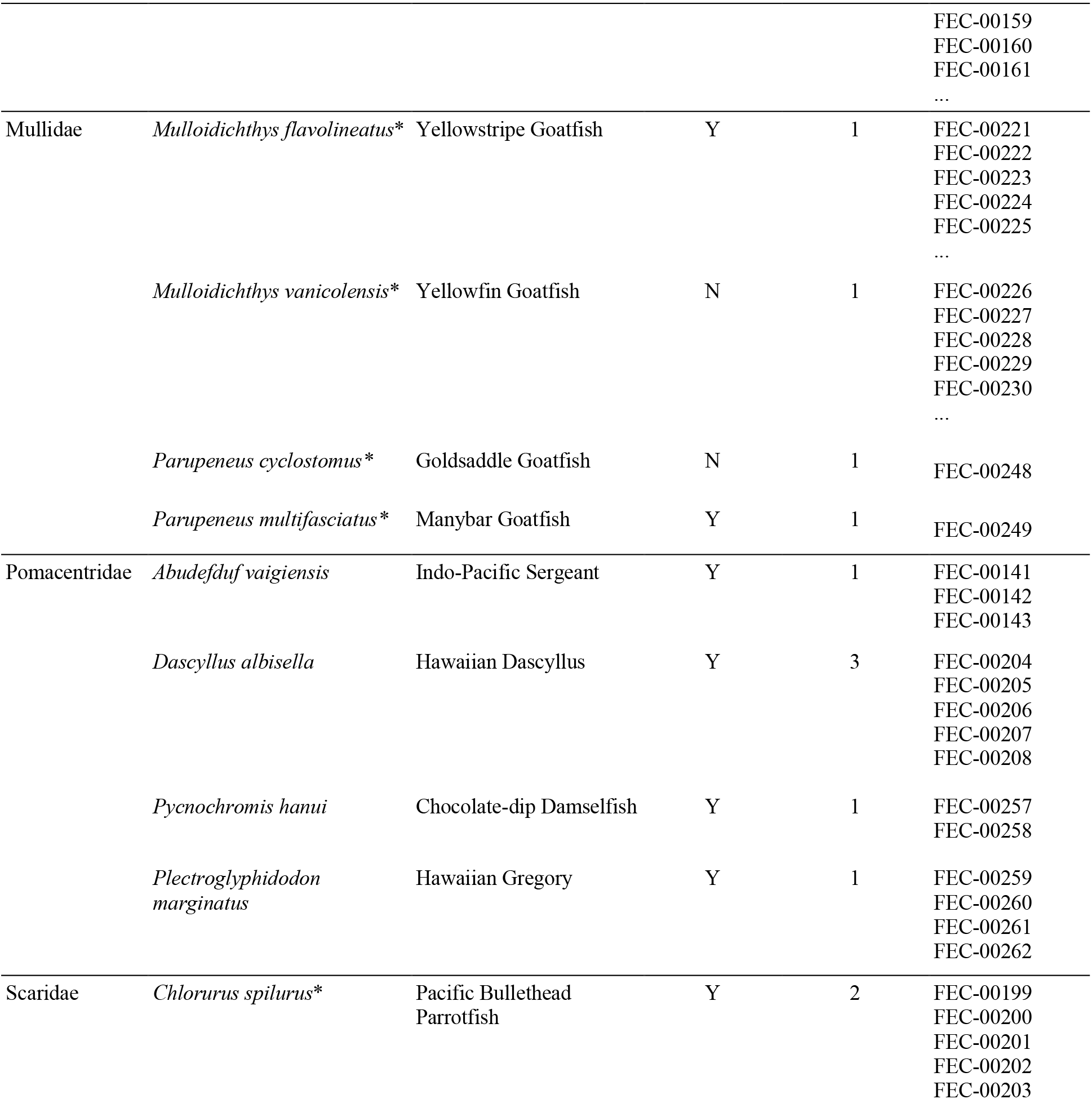
Identified sounds from 31 species of Hawaiian reef fishes. Representative specimens for each species are available at fisheyecollaborative.org/library. In cases where more than five video-validated sounds were recorded for a species, indicated by ellipses following catalog numbers, additional recordings are available upon request. Commercial species (per Hawaii DLNR) are indicated with an asterisk. Species names follow the taxonomy in Eschmeyer’s Catalog of Fishes (Fricke et al. 2026).

## Results

With this method, we identified sounds from 31 species across 14 families (Table 1). Of these species, 13 had never been previously documented in the literature to produce sounds (Looby et al. 2026). For the 18 species that we recorded that had been previously documented as sound-producing, all were included in the foundational study by Tricas and Boyle (2014). According to FishSounds.net, only one species, *M. berndti,* has independently peer-reviewed published audio recordings. Salmon (1967) published both spectrograms and recordings for *M. berndti* which are supported by our recordings of that species. *D. albisella* has been the subject of multiple studies (Lobel and Mann 1995; Mann and Lobel 1995; Mann and Lobel 1997; Mann and Lobel 1998) and many examples of their sounds appear online, including on YouTube, there are no sounds published as part of the scientific studies. One recording from SanctSound that is labeled “damselfish” is likely *D. albisella* (see https://sanctsound.ioos.us/sounds.html#damselfish-papahnaumokukea).

The reframed video and soundfield visualizations (Fig. 3) allowed for identification of species-specific sounds associated with behaviors such as courtship (Fig. 3A, 3B) and agonistic interactions (Fig. 3C). Other distinctive sounds emerged including vocalizations with unknown behavioral contexts (Fig. 3D, 3E) and sounds associated with feeding (Fig. 3F). Some species produced sounds but were less distinct from one another (Fig. 3G, 3H).

The intention of this study is not just to identify sounds but to identify opportunities for acoustic categorizations that could be used for monitoring. Many fish sounds are very similar, however, for some taxa, identified in Table 1, sounds are easily distinguishable by trained observers. The courtship sounds of *D. albisella* and *S. bursa* were among the most common fish sounds throughout our deployments and can be readily identified to the species level. Similar sounds from congeners *M. berndti* and *M. kuntee* can be ascribed to the genus *Myripristis.* Various sounds from four wrasse species (*A. chrysocephalus*, *B. albotaeniatus*, *P. evanidus*, and *T. duperrey*) were recorded with unclear behavioral contexts (Fig. 3D, 3E) and shared many characteristics. These wrasse sounds are qualitatively human-identifiable to the family level (Labridae) for this study, but more extensive data may allow them to be differentiated further. Corroborating the human qualitative identifiability scores, feeding sounds (N=58) and communication sounds (N= 394) were significantly different in maximum entropy (Kruskal-Wallis: χ^2^ = 94.8892, DF = 1, P< 0.0001).

The DFA based on sound measurements from our focal taxonomic groups showed a classification consistent with human identifiability (Table 1). The first canonical discriminant functions explained 86.43% of the variance, with canonical structure showing that canonical variable 1 explaining 50.23% of the variance (primarily influenced by Center Frequency, Delta Frequency, and Bandwidth 90%), and canonical variable 2 explained 36.20 of the variance (primarily influenced by Aggregate Entropy, Average Entropy, and Maximum Entropy) (Fig. 3, canonical structure variables can be seen in Table S2). The DFA correctly classified sounds from the four taxa at >87.5% (Table 2). The DFA showed a low misclassification rate across all groups at < 12.5%. The high correct classification scores and low misclassification scores suggest that these 12 acoustic parameters can successfully differentiate sounds between these groups (Wilks’ Lambda = 0.0246, F=48.937, DF= 48, 1454.3, *p* < 0.0001).

**Table 2.** Classification performance of DFA on acoustic measurements for the different acoustic functional groups (from Fig. 3), showing classification rates. Sound measurements are described in Table S1. Cells are shaded proportionately on a scale of white (0) to black (1).

|  |  | Predicted Group |  |  |  |  |
| --- | --- | --- | --- | --- | --- | --- |
|  |  | *Dascyllus albisella* | *Sufflamen bursa* | *Myripristis* sp. | Labridae | Other |
| Actual | *Dascyllus albisella* | **0.903** | 0.000 | 0.000 | 0.011 | 0.086 |
|  | *Sufflamen bursa* | 0.000 | **0.985** | 0.000 | 0.007 | 0.007 |
|  | *Myripristis* sp. | 0.000 | 0.000 | **0.875** | 0.000 | 0.125 |
|  | Labridae | 0.000 | 0.000 | 0.000 | **0.970** | 0.030 |
|  | Other | 0.012 | 0.012 | 0.000 | 0.037 | **0.939** |

These results combine to suggest that sounds from these taxa are reliably separable both manually and statistically. While some overlap exists between classes (Fig. 5), the DFA shows separation between classes even with a limited dataset and relatively few acoustic measurements. As a proof-of-concept of identifying these sound classes in PAM data, we analyzed the temporal occurrence of fish sounds from a continuous 30 min audio-only PAM-equivalent recording (Fig. 5). In addition to these 4 vocalization sound classes, we were able to identify feeding sounds from the two species of grazers we recorded, *C. dumerilii* and *C. spilurus.* These sounds cannot yet be distinguished from each other but can be ascribed to a single functional group, ‘grazers’.

## Discussion

Our 31-species Hawaiian fish sound library represents the largest open-access collection of *in situ* Pacific fish sound recordings in existence to date and represents the first documented sound production for 13 species. For 17 other species, our recordings are the first field-based scientific recordings publicly available. Because the 360° method captures entire behavioral sequences, for species with more extensive documentation of natural acoustic behaviors, such as *D. albisella* (e.g., Lobel and Mann 1995), *M. berndti* (Salmon 1967), and *S. bursa* (Salmon et al. 1968), our library now contains video and acoustic recordings that can help interpret repertoires by adding further behavioral context. Our study builds on the 49 Caribbean species already documented using UPAC-360 systems, contributing a substantial taxonomic and geographic expansion to the open-access library (Dantzker et al. 2025c). Our results further demonstrate that fishes across an increasing range of taxa not only produce sounds, but that many produce repeated, distinctive, and identifiable acoustic signals which may in the future be used for valuable ecosystem monitoring applications.

### Identifying sounds in PAM datasets

Many of the species we recorded produced pulsed sounds that at an initial glance may appear similar to one another. Despite these superficial similarities, DFA on selected sound measurements from Raven (Fig. 4) revealed clear separation among several species, families, and functional groups. This separation emerged despite the use of a relatively small number of acoustic metrics, including temporal and spectral parameters. Among the species that produced single pulses, we were able to identify those made by *S. bursa* and those made by wrasses (Labridae) through manual spectrogram review of PAM data (Fig. 5), and our DFA results reflect the separation of those groups from other species (Fig. 4). Among the species that produced multipulsed calls, we manually identified *D. albisella, Myripristis spp.,* and additional labrid species, further supported by the separation in Fig. 4. While broadband grazing sounds from herbivory have been widely described and are frequently identified aurally (Sartori and Bright 1973; Tricas and Boyle 2021), we have found that, in addition, a trained expert using the video-verified species-attributed examples from this study can hear differences in feeding sounds across taxa. For example, we were able to distinguish algal grazing sounds from those related to feeding on invertebrates.

**Figure 4.**
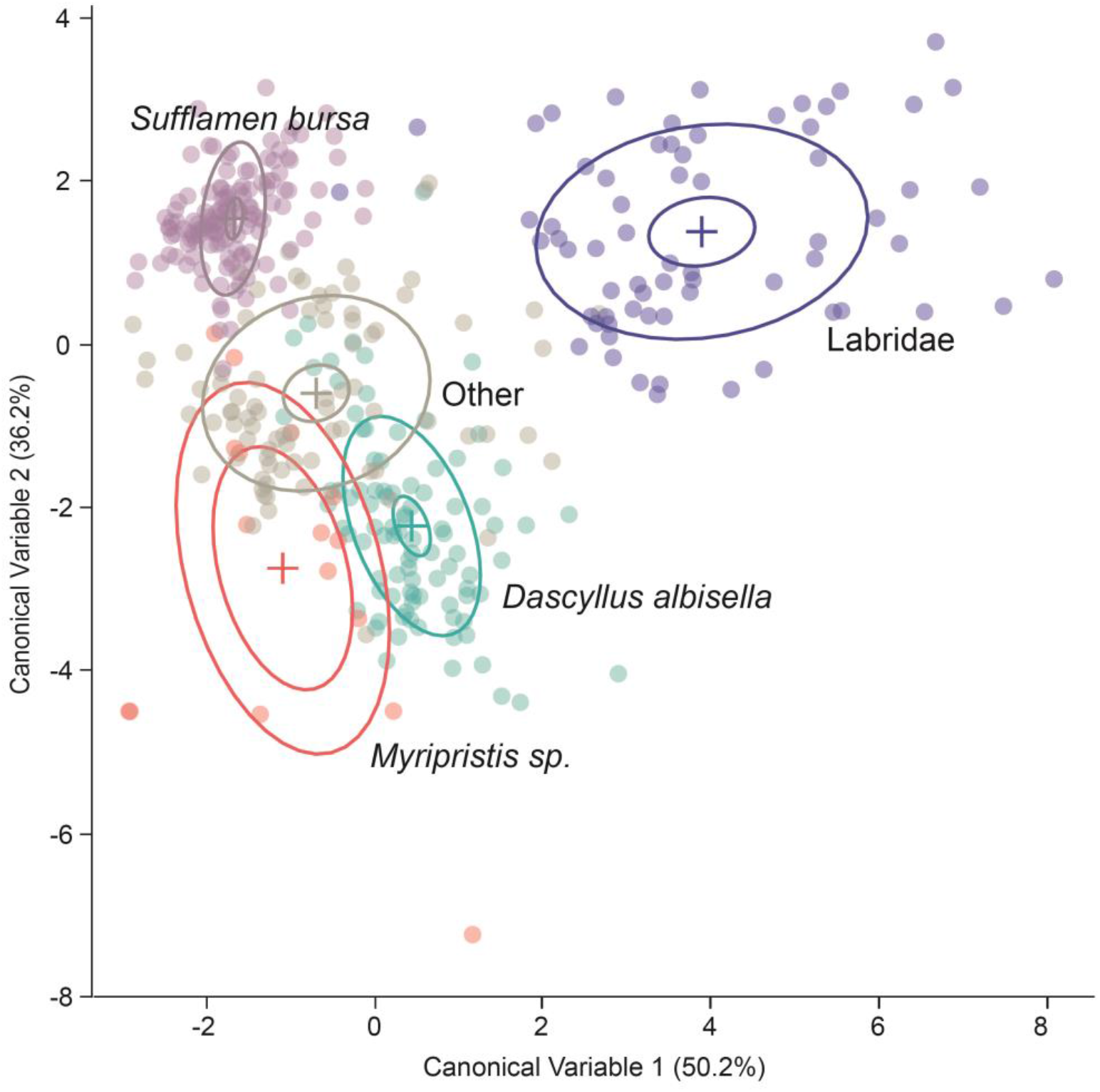
DFA of sound measurements from *D. albisella*, *S. bursa*, *Myripristis* sp., Labridae, and all other species. DFA shows the first two canonical discriminant functions, and the ellipses contain 95% of the points for that class.

**Figure 5.**
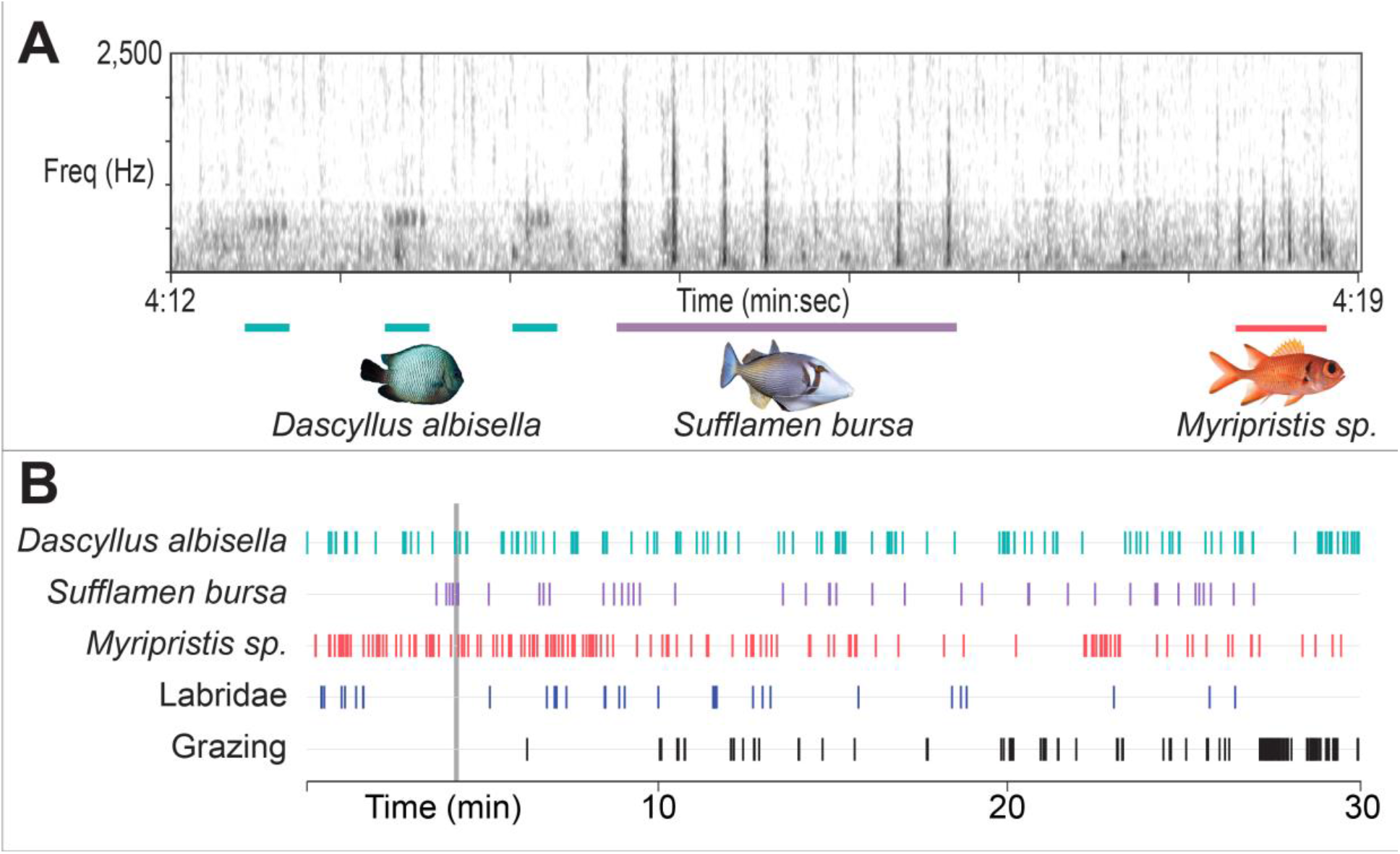
A) A seven-second spectrogram taken from the PAM recording with representative sounds from three identifiable taxa. Colored bars on x axis highlight the duration of the call of the underlying species. B) Raster plot of temporal occurrences of identifiable sounds over the duration of a 30-minute recording (spectrogram not shown). Fish photos by ANR.

It is likely that a more comprehensive analysis with additional metrics or feature embeddings (Best et al. 2023; Ghani et al. 2023) could yield greater separation between classes and would likely be more predictive of how machine learning algorithms would perform for automated detection and classification (Duane et al. 2026; Williams et al. 2025) needed for identifying sounds in large PAM datasets. Now, with the ability to make these open access, others can identify and differentiate sounds from more fish species. These validated sounds can be used for cross-comparison of signals in other PAM recordings or to develop traditional and AI detectors and classifiers for Hawaiian and Pacific reef fishes.

There were some sounds that stood out as distinct and identifiable that were not captured in Fig. 2. For example, goatfishes in the family Mullidae produced unique low-frequency rumbles when startled. Some sounds produced by labrid species are very different from one another and can be tentatively identified to a species level. Due to the relatively few examples attributed to these taxa, it is not yet clear whether more data would allow us to reliably distinguish these sounds. The taxa included in our PAM analysis represent the most consistently vocal species for which we have ample data to back confident acoustic identifications. With ongoing data collection, we could likely begin to identify additional species in PAM recordings, including commercially important taxa like goatfishes.

### Acoustic indicator species

A broader goal in using PAM to understand coral reef ecology and inform conservation and management is to use species-specific acoustic signals as ways to infer broader ecological processes. The acoustically distinct taxa (levels 2 and 3 in Table 1) produce sounds that are consistent, detectable, and behaviorally informative, providing valuable insights into the health of Hawaiian reefs. For example, the endemic Hawaiian Dascyllus (*D. albisella*) emerged as one of the most acoustically distinctive, and well-studied (Laboury et al. 2025; Lobel and Mann 1995; Mann and Lobel 1995; Mann and Lobel 1997; Mann and Lobel 1998) taxa we recorded and is a strong candidate for machine learning detectors, as it emerges as a distinct class in cluster analyses (Duane et al. 2026). *Dascyllus* are dependent on live coral structure across their life stages, establishing territories and nests on coral heads upon recruitment (Danilowicz 1995; Stevenson 1963). Detection of *Dascyllus* territorial and reproductive acoustic activity can reflect habitat occupancy, reef structural integrity, and habitat connectivity.

Lei Triggerfish (*S. bursa*) produced frequent sounds during extended courtship interactions, and both soldierfish species (*M. berndti* and *M. kuntee*) produced distinct pulse trains during agonistic interactions. These species are subject to localized fishing pressure, and changes in acoustic detections can reflect the efficacy of protected areas and no-take zones. Additionally, *Myripristis* alarm calls may also indicate the occurrence of nearby predators (Popper et al. 1973). Herbivorous grazing fishes such as *C. spilurus*, another commercially harvested species, are among the most functionally important groups on coral reefs, suppressing macroalgal growth and supporting coral recruitment and reef resilience (Bonaldo et al. 2014; Tricas and Boyle 2021). Acoustic detection of grazing can provide early warning signs of declining herbivory threatening reef integrity.

### Uses for low identifiability species sounds

For the many species we typed with an identifiability of 1, it is not yet possible to identify their sounds in raw PAM recordings. Additional sampling will likely increase the number of attributable sounds, allowing for identification of important signal features that further support their species-specificity. Further study may reveal other sounds in their repertoires, perhaps associated with mating or territoriality, or other social behaviors involving sounds (van Oosterom et al. 2016). However, assigning even the least distinctive sounds to species is important, as it provides better inference into the relationship between acoustic diversity and species diversity on coral reefs. It allows greater certainty that the distinctive sounds you are assigning are unique if you know the sound from other species that are present in the habitat. And, when training AI detectors, sound verified to have come from other species can serve as “true negatives.” Documenting these less identifiable sounds also points at the danger of using subjective grouping based on spectrograms to classify unknown sounds into classes. Having attributed sounds to so many species, we can confidently say that superficially similar sounds can come from widely divergent taxa.

### Inferring functional diversity

Coral reef resilience depends not only on species richness, but on the maintenance of functional diversity and key ecosystem processes (Bellwood et al. 2004; Brandl et al. 2019; Mouillot et al. 2014) including algal suppression, trophic transfer, nutrient cycling, and benthic community structure (Brandl et al. 2019; Bellwood et al. 2019). Hawaiian reefs may be particularly vulnerable to losses in functional diversity due to their relatively low species richness and high levels of endemism compared to Indo-Pacific biodiversity hotspots (Allen 2008; DeMartini and Friedlander 2004; Kulbicki et al. 2013; Lobel et al. 2020). Although we cannot yet identify all of our recorded species with precise taxonomic resolution, our identifiable groups serve unique functional roles (sensu Wolfe et al. 2020), and their acoustic activity can provide a snapshot of the functional diversity of the reef (Jarriel et al. 2024; Tricas and Boyle 2021). The functional groups represented include site-attached planktivores (*D. albisella*) that provide localized nutrient cycling, nocturnal planktivores (*Myripristis spp.*) that transfer pelagic energy to reefs, mobile benthic invertivores (*S. bursa*) that serve as both predators and prey for larger fishes, and grazers and bioeroders (*C. spilurus* and *C. dumerilii*). At the functional-acoustic level for our analysis, here “wrasses” (Labridae) includes *A. chrysocephalus*, *B. albotaeniatus*, *P. evanidus*, and *T. duperrey* that feed on small benthic invertebrates. Because ecological functions are not evenly distributed across reef fish assemblages, changes in the acoustic activity of key taxa may correspond to disproportionate ecological consequences (Bellwood et al. 2012; Mouillot et al. 2013). Application of our results in PAM therefore has the potential to assess not only species presence, but broader patterns of ecosystem function, ecological stability, and reef resilience.

### Application in Pacific reefs

This study was conducted in one of Hawai’i’s many Marine Protected Areas (MPAs) (Fig. 1). Of the species recorded in this study, approximately half (16) are part of commercial or subsistence fisheries and are harvested on Hawaiian reefs (Table 1; see https://dlnr.ehawaii.gov/cmls-fr/pdf/SpeciesList.pdf). The establishment of this validated fish sound library opens the doors for widespread application of PAM techniques in Hawaiian MPAs, supplementing or partially replacing existing monitoring methods such as visual census, BRUVs, and eDNA that, when used alone, may yield incomplete biodiversity data (Apprill et al. 2023; Kotwicki et al. 2026; Madin et al. 2019; Mooney et al. 2020; Muenzel et al. 2024). PAM is relatively low-cost and ideal for assessing long-term trends, detecting species shifts due to climate change and overfishing, monitoring protected area compliance, and may provide early warnings of declines in fish populations. The use of PAM as a monitoring tool will become increasingly scalable with the development of detection algorithms that greatly reduce the need for manual annotation of sounds (Mooney et al. 2020).

### Limitations and future work

Data collection for this study was conducted exclusively during the daytime at a single site over three days. The constraints in spatial and temporal coverage and minimal variation in light, depth, and temperature resulted in a limited glimpse into the soundscape of Hawaiian reefs. Further deployments across varying times of day, seasons, depths, environmental gradients, and benthic compositions can greatly expand our library and our understanding of the acoustic repertoires of fishes, particularly for uncommon or infrequently vocalizing species. Fish are particularly acoustically active at dawn and dusk (Azofeifa-Solano et al. 2025; La Manna et al. 2024; McCauley and Cato 2000; Raick et al. 2023), and this study did not encompass nocturnal activity. However, supplemental light sources could be applied to capture sounds from crepuscular and nocturnal fishes, as well as cryptic species that make sounds from inside caves and crevices during the day but might vocalize in the open at other times (Dantzker et al. 2025b). Expanding the geographic range of deployment sites will provide a better understanding of regional call variability (Mann and Lobel 1998; Parmentier et al. 2005) and the transferability of results to a large-scale acoustic monitoring network across Pacific reefs.

## Conclusions

This study substantially expands the known acoustic diversity of Hawaiian reef fishes through the development of the largest open-access in-situ Pacific reef fish sound library currently available. Our findings demonstrate that many Hawaiian reef fishes produce repeated, distinctive, and ecologically informative acoustic signals detectable under realistic PAM conditions. Preliminary DFA results further suggest that multiple species and functional groups can be reliably classified, supporting the feasibility of species- and guild-level acoustic monitoring on coral reefs. The acoustically identifiable taxa represent several ecologically important reef functions, including planktivory, nutrient cycling, benthic predation, and herbivory. They further serve as indicators of habitat complexity, coral health, and fisheries activity. These applications may be particularly valuable across Pacific coral reefs, where many ecosystems remain remote, under-monitored, and increasingly vulnerable to climate change, overfishing, and habitat degradation. By linking species-specific acoustic activity to reef ecological processes, this work helps bridge the gap between emerging acoustic technologies and actionable biodiversity conservation and fisheries management. Continued expansion of acoustic reference libraries and automated detection workflows may ultimately allow reef soundscapes to serve as scalable indicators of coral reef condition across the Pacific.

## Statements and Declarations

### Author Contributions

**Erika Berlik:** Data Curation, Formal Analysis, Investigation, Methodology, Visualization, Writing – Original Draft Preparation; Writing – Review & Editing; **Marc Dantzker:** Conceptualization, Data Curation, Field Work, Formal Analysis, Funding Acquisition, Investigation, Methodology, Project Administration, Supervision, Visualization, Writing – Original Draft Preparation, Writing – Review & Editing; **Symeon Delikaris-Manias:** Field work, Investigation, Methodology, Writing – Review & Editing; **Matthew Duggan:** Methodology, Writing – Review & Editing; **Aaron Rice:** Conceptualization, Field Work, Formal Analysis, Funding Acquisition, Investigation, Methodology, Visualization, Writing – Original Draft Preparation, Writing – Review & Editing

### Data Availability

Our fish sound specimens are available via our library at fisheyecollaborative.org/library, as well as via FishEye Collaborative’s data in the Global Biodiversity Information Facility (GBIF) and the Ocean Biodiversity Information System (OBIS).

## Supplemental Material

**Table S1.**
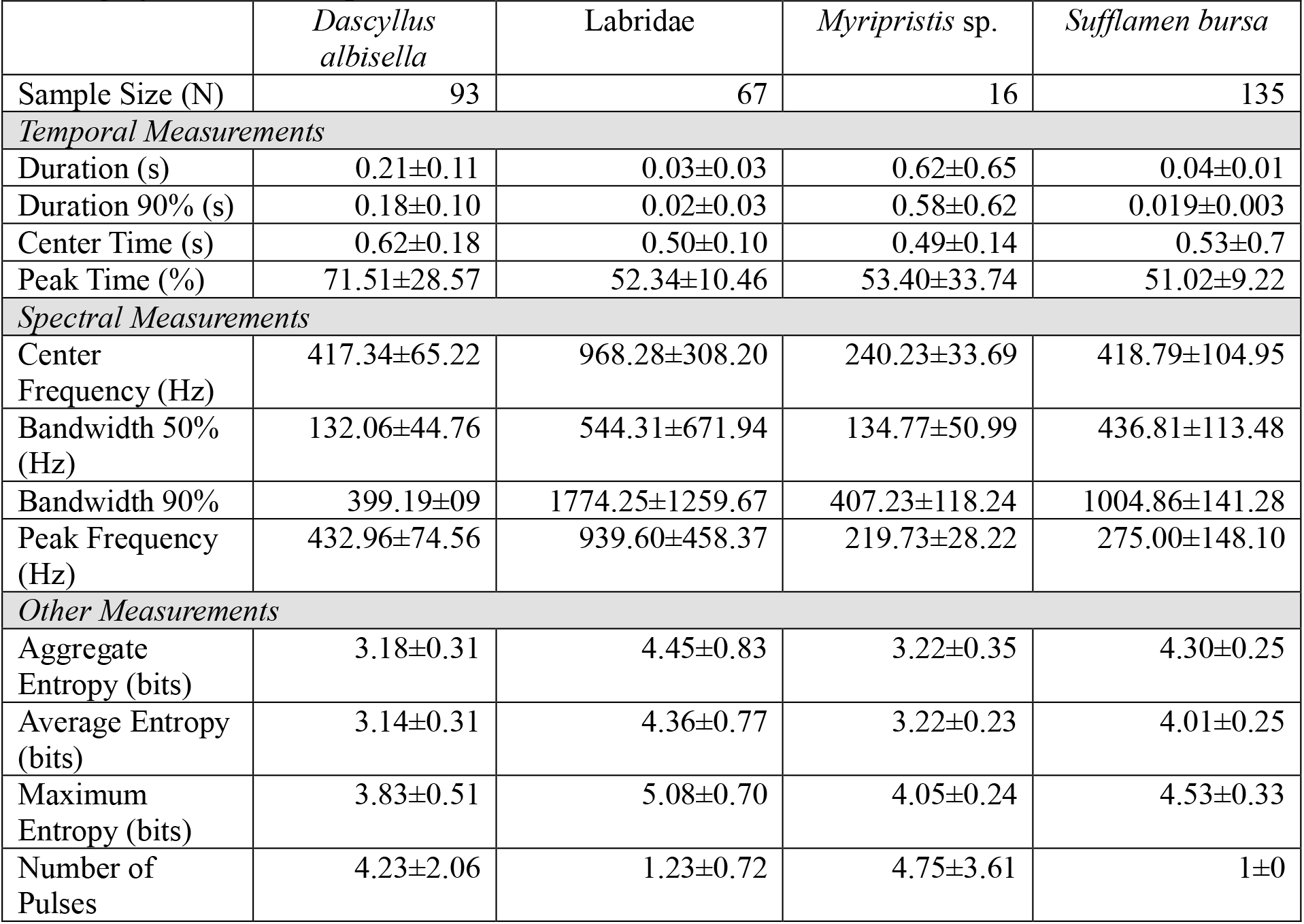
Signal measurements for different focal taxonomic groups, measured in Raven Pro 1.6.5. Values are represented as mean ± standard deviation. Note: For *S. bursa*, all pulses were counted individually despite occurring in quick succession. This is because pulses were not linked in to one another in discrete, stereotyped, structured “calls,” but rather occurred in inconsistent bursts with highly variable interpulse intervals.

**Table S2.** Canonical structure from Discriminant Function Analysis with signal properties included in analysis.

|  | **Canonical 1** | **Canonical 2** | **Canonical 3** | **Canonical 4** |
| --- | --- | --- | --- | --- |
| **Bandwidth 50%** | 0.158037 | 0.540656 | 0.168781 | 0.086838 |
| **Bandwidth 90%** | 0.440593 | 0.62105 | 0.225291 | 0.041796 |
| **Aggregate Entropy** | 0.136329 | 0.846783 | 0.272564 | 0.171342 |
| **Average Entropy** | 0.261644 | 0.786229 | 0.34547 | 0.194121 |
| **Maximum Entropy** | 0.360332 | 0.665978 | 0.478027 | 0.256204 |
| **Center Time** | 0.029037 | -0.23893 | 0.117051 | 0.7397 |
| **Center Frequency** | 0.803818 | 0.400824 | -0.01386 | -0.02825 |
| **Duration** | -0.05575 | -0.59895 | 0.702433 | -0.11256 |
| **Duration 90%** | -0.03952 | -0.58414 | 0.711924 | -0.15633 |
| **Peak Frequency** | 0.785862 | 0.134638 | -0.11642 | -0.1232 |
| **Peak Time** | 0.106658 | -0.35413 | 0.107016 | 0.651454 |
| **Pulses** | 0.068635 | -0.76587 | 0.268222 | 0.315395 |

## References

Allen GR (2008) Conservation hotspots of biodiversity and endemism for Indo-Pacific coral reef fishes. Aquat Conserv Mar Freshw Ecosyst 18:541–556 10.1002/aqc.880

Andrello M, Darling ES, Wenger A, Suarez-Castro AF, Gelfand S, Ahmadia GN (2022) A global map of human pressures on tropical coral reefs. Conserv Lett 15:e12858 10.1111/conl.12858

Apprill A et al. (2023) Toward a new era of coral reef monitoring. Environ Sci Technol 57:5117–5124 10.1021/acs.est.2c05369

Au WWL, Mobley J, Burgess WC, Lammers MO, Nachtigall PE (2000) Seasonal and diurnal trends of chorusing humpback whales wintering in waters off western Maui. Mar Mamm Sci 16:530–544 10.1111/j.1748-7692.2000.tb00949.x

Au WWL, Richlen M, Lammers MO (2012) Soundscape of a nearshore coral reef near an urban center. In: Popper AN, Hawkins A (eds) The Effects of Noise on Aquatic Life, vol 730. Advances in Experimental Medicine and Biology. Springer New York, pp 345–351. 10.1007/978-1-4419-7311-5_78

Azofeifa-Solano JC et al. (2025) Soundscape analysis reveals fine ecological differences among coral reef habitats. Ecol Indic 171:113120 10.1016/j.ecolind.2025.113120

Bellwood DR, Hoey AS, Hughes TP (2012) Human activity selectively impacts the ecosystem roles of parrotfishes on coral reefs. Proc R Soc B 279:1621–1629 10.1098/rspb.2011.1906

Bellwood DR, Hughes TP, Folke C, Nystrom M (2004) Confronting the coral reef crisis. Nature 429:827–833 10.1038/nature02691

Bertucci F, Parmentier E, Lecellier G, Hawkins AD, Lecchini D (2016) Acoustic indices provide information on the status of coral reefs: an example from Moorea Island in the South Pacific. Sci Rep 6:33326 10.1038/srep33326

Best P, Paris S, Glotin H, Marxer R (2023) Deep audio embeddings for vocalisation clustering. PLOS ONE 18:e0283396 10.1371/journal.pone.0283396

Boehlert GW (1993) Fisheries and marine resources of Hawaii and the U.S.-associated Pacific Islands: An introduction. Mar Fish Rev 55:3–6

Bonaldo RM, Hoey AS, Bellwood DR (2014) The ecosystem roles of parrotfishes on tropical reefs. Oceanogr Mar Biol Ann Rev 52:81–132 10.1201/b17143-3

Brandl SJ, Rasher DB, Côté IM, Casey JM, Darling ES, Lefcheck JS, Duffy JE (2019) Coral reef ecosystem functioning: eight core processes and the role of biodiversity. Front Ecol Environ 17:445–454 10.1002/fee.2088

Carrillo-García DM, Kolb M (2022) Indicator Framework for Monitoring Ecosystem Integrity of Coral Reefs in the Western Caribbean. Ocean Sci J 57:1–24 10.1007/s12601-022-00055-1

Cornell K. Lisa Yang Center for Conservation Bioacoustics (KLY-CCB) (2022) Raven Pro: Interactive Sound Anaysis Software (Version 1.6.5) Cornell Lab of Ornithology, Ithaca, NY. Available at: https://ravensoundsoftware.com/

Costa BM, Kendall MS (eds) (2016) Marine Biogeographic Assessment of the Main Hawaiian Islands. NOAA Technical Memorandum NOS NCCOOS 214. National Centers for Coastal Ocean Science, National Oceanic and Atmospheric Administration, Silver Spring, MD. 10.7289/V5/TM-NOS-NCCOS-214

Danilowicz BS (1995) Spatial patterns of spawning in the coral reef damselfish *Dascyllus albisella*. Mar Biol 122:145–155 10.1007/bf00349288

Dantzker MS, Duggan MT, Berlik E, Delikaris-Manias S, Bountourakis V, Pulkki V, Rice AN (2025a) Deciphering complex coral reef soundscapes with spatial audio and 360° video. Methods Ecol Evol 16:2622–2637 10.1111/2041-210x.70149

Dantzker MS et al. (2025b) Opening coral reef soundscapes to species-level monitoring by identifying fish sounds. In: IEEE OCEANS 2025 - Great Lakes, 29 Sept.–2 Oct. 2025. pp 1–10. 10.23919/OCEANS59106.2025.11245117

Dantzker MS, Berlik E, Duggan MD, Delikaris-Manias S, Rice AN (2025c) FishEye Collaborative Fish Sound Library. FishEye Collaborative. https://www.fisheyecollaborative.org/library. Accessed 20 July 2026

Davidson IK, Williams B, Stratford JE, Chapuis L, Simpson SD, Radford AN (2024) Context-dependent multimodal behaviour in a coral reef fish. R Soc Open Sci 11:240151 doi:10.1098/rsos.240151

Delikaris-Manias S, McCormack L, Huhtakallio I, Pulkki V (2018) Real-time underwater spatial audio: A feasibility study. Proceedings of the 144th Convention of the Audio Engineering Society. Milan, Italy

Delikaris-Manias S, Pulkki V (2013) Cross pattern coherence algorithm for spatial filtering applications utilizing microphone arrays. IEEE Trans Audio Speech Lang Process 21:2356–2367 10.1109/TASL.2013.2277928

DeMartini EE, Friedlander AM (2004) Spatial patterns of endemism in shallow-water reef fish populations of the Northwestern Hawaiian Islands. Mar Ecol Prog Ser 271:281–296 10.3354/meps271281

Duane D, Dolan K, Iafrate J, Freeman L The effect of COVID-19 shutdowns on Hawaiian coral reef soundscapes. In: IEEE OCEANS 2023 - Limerick, 5–8 June 2023 2023. pp 1–6. 10.1109/OCEANSLimerick52467.2023.10244572

Duane D, Duggan MT, Berlik E, Dantzker MS, Rice AN, Freeman L (2026) Multi-class, unsupervised detection and classification of biological and anthropogenic sounds in coral reefs. PLoS Comput Biol 22:e1014516 10.1371/journal.pcbi.1014516

Duane D, Freeman S, Freeman L (2024) Moonlight-driven biological choruses in Hawaiian coral reefs. PLOS ONE 19:e0299916 10.1371/journal.pone.0299916

Edmunds PJ (2024) Why keep monitoring coral reefs? BioScience 74:552–560 10.1093/biosci/biae046

Ferrier-Pagès C, Leal MC, Calado R, Schmid DW, Bertucci F, Lecchini D, Allemand D (2021) Noise pollution on coral reefs? — A yet underestimated threat to coral reef communities. Mar Pollut Bull 165:112129 10.1016/j.marpolbul.2021.112129

Flower J et al. (2017) Interpreting coral reef monitoring data: A guide for improved management decisions. Ecol Indic 72:848–869 10.1016/j.ecolind.2016.09.003

Freeman LA, Freeman SE (2016) Rapidly obtained ecosystem indicators from coral reef soundscapes. Mar Ecol Prog Ser 561:69–82 10.3354/meps11938

Freeman SE, Rohwer FL, D’Spain GL, Friedlander AM, Gregg AK, Sandin SA, Buckingham MJ (2014) The origins of ambient biological sound from coral reef ecosystems in the Line Islands Archipelago. J Acoust Soc Am 135:1775–1788 10.1121/1.4865922

Fricke R, Eschmeyer WN, Fong JD (2026) Eschmeyer’s Catalog of Fishes: Genera, Species, References. Electronic version accessed 1 June 2026. Available at: http://researcharchive.calacademy.org/research/ichthyology/catalog/fishcatmain.asp,

Friedlander A et al. (2008) The state of coral reef ecosystems of the main Hawaiian Islands. NOAA Tech Mem NOS NCCOS 73:219–264

Ghani B, Denton T, Kahl S, Klinck H (2023) Global birdsong embeddings enable superior transfer learning for bioacoustic classification. Sci Rep 13:22876 10.1038/s41598-023-49989-z

Greenfield DW (2003) A survey of the small reef fishes of Kane’ohe Bay, O’ahu, Hawaiian Islands. Pac Sci 57:45–76 10.1353/psc.2003.0001

Harris SA, Shears NT, Radford CA (2016) Ecoacoustic indices as proxies for biodiversity on temperate reefs. Methods Ecol Evol 7:713–724 10.1111/2041-210X.12527

Haver SM et al. (2018) Monitoring long-term soundscape trends in U.S. Waters: The NOAA/NPS Ocean Noise Reference Station Network. Mar Policy 90:6–13 10.1016/j.marpol.2018.01.023

Havlik M-N, Predragovic M, Duarte CM (2022) State of play in marine soundscape assessments. Front Mar Sci 9:919418 10.3389/fmars.2022.919418

Heenehan HL, Van Parijs SM, Bejder L, Tyne JA, Southall BL, Southall H, Johnston DW (2017) Natural and anthropogenic events influence the soundscapes of four bays on Hawaii Island. Mar Pollut Bull 124:9–20 10.1016/j.marpolbul.2017.06.065

Hodson EJ, Cox K, Juanes F, Looby A (2025) Actively soniferous tropical reef fishes are diverse, vulnerable, and valuable. J Fish Biol 106:990–995 10.1111/jfb.16030

Hughes TP et al. (2017) Coral reefs in the Anthropocene. Nature 546:82–90 10.1038/nature22901

Ibrahim AK, Zhuang HQ, Schaerer-Umpierre M, Woodward C, Erdol N, Cherubin LM (2024) Fish Acoustic Detection Algorithm Research: a deep learning app for Caribbean grouper calls detection and call types classification. Front Mar Sci 11:1378159 10.3389/fmars.2024.1378159

Jarriel SD, Formel N, Ferguson SR, Jensen FH, Apprill A, Mooney TA (2024) Unidentified fish sounds as indicators of coral reef health and comparison to other acoustic methods. Front Remote Sens 5:1338586 10.3389/frsen.2024.1338586

Jokiel PL (2008) Biology and Ecological Functioning of Coral Reefs in the Main Hawaiian Islands. In: Riegl BM, Dodge RE (eds) Coral Reefs of the USA. Springer Netherlands, Dordrecht, pp 489–517. 10.1007/978-1-4020-6847-8_12

Kaplan MB, Lammers MO, Zang E, Mooney TA (2018) Acoustic and biological trends on coral reefs off Maui, Hawaii. Coral Reefs 37:121–133 10.1007/s00338-017-1638-x

Kaplan MB, Mooney TA, Partan J, Solow AR (2015) Coral reef species assemblages are associated with ambient soundscapes. Mar Ecol Prog Ser 533:93–107 10.3354/meps11382

Kotwicki S et al. (2026) Pathways toward modernizing fisheries-independent surveys. ICES J Mar Sci 83:fsag134 10.1093/icesjms/fsag134

Kügler A, Lammers MO, Pack AA, Tenorio-Hallé L, Thode AM (2024) Diel spatio-temporal patterns of humpback whale singing on a high-density breeding ground. R Soc Open Sci 11:230279 doi:10.1098/rsos.230279

Kulbicki M et al. (2013) Global biogeography of reef fishes: A hierarchical quantitative delineation of regions. PLoS ONE 8:e81847 10.1371/journal.pone.0081847

La Manna G, Moro Merella M, Vargiu R, Morello G, Sarà G, Ceccherelli G (2024) Spatio-temporal patterns of fish acoustic communities in Western Mediterranean coralligenous reefs: optimizing monitoring through recording duration. Front Mar Sci 11:1483661 10.3389/fmars.2024.1483661

Laboury S, Parmentier E, Lobel PS (2025) Are there individual acoustic signatures in the damselfish *Dascyllus albisella*? J Acoust Soc Am 157:48–56 10.1121/10.0034790

Lammers MO et al. (2023) The occurrence of humpback whales across the Hawaiian archipelago revealed by fixed and mobile acoustic monitoring. Front Mar Sci 10:1083583 10.3389/fmars.2023.1083583

Lin T-H, Akamatsu T, Sinniger F, Harii S (2021) Exploring coral reef biodiversity via underwater soundscapes. Biol Conserv 253:108901 10.1016/j.biocon.2020.108901

Lindseth A, Lobel P (2018) Underwater soundscape monitoring and fish bioacoustics: A review. Fishes 3:36 10.3390/fishes3030036

Lobel PS, Kaatz IM, Rice AN (2010) Acoustical behavior of coral reef fishes. In: Cole KS (ed) Reproduction and Sexuality in Marine Fishes: Evolutionary Patterns and Innovations. University of California Press, Berkeley, CA, pp 307–386. 10.1525/california/9780520264335.003.0010

Lobel PS, Lobel LK, Randall JE (2020) Johnston Atoll: Reef Fish Hybrid Zone between Hawaii and the Equatorial Pacific. Diversity 12:83 10.3390/d12020083

Lobel PS, Mann DA (1995) Spawning sounds of the damselfish, *Dascyllus albisella* (Pomacentridae), and relationship to male size. Bioacoustics 6:187–198 10.1080/09524622.1995.9753289

Looby A et al. (2023) Global inventory of species categorized by known underwater sonifery. Sci Data 10:892 10.1038/s41597-023-02745-4

Looby A et al. (2026) FishSounds. http://www.fishsounds.net, version 2026-Apr. Accessed 20 July 2026.

MacNeil MA et al. (2015) Recovery potential of the world’s coral reef fishes. Nature 520:341– 344 10.1038/nature14358

Madin EMP, Darling ES, Hardt MJ (2019) Emerging technologies and coral reef conservation: Opportunities, challenges, and moving forward. Front Mar Sci 6:00727 10.3389/fmars.2019.00727

Madrigal BC, Kügler A, Zang EJ, Lammers MO, Hatch LT, Pacini AF (2024) Comparing the underwater soundscape of the Hawaiian Islands Humpback Whale National Marine Sanctuary and potential influences of the COVID-19 pandemic. Front Mar Sci Volume 11–2024 10.3389/fmars.2024.1342454

Mahale VP, Chanda K, Chakraborty B, Salkar T, Sreekanth GB (2023) Biodiversity assessment using passive acoustic recordings from off-reef location—Unsupervised learning to classify fish vocalization. J Acoust Soc Am 153:1534–1553 10.1121/10.0017248

Mann DA, Lobel PS (1995) Passive acoustic detection of sounds produced by the damselfish, *Dascyllus albisella* (Pomacentridae). Bioacoustics 6:199–213 10.1080/09524622.1995.9753290

Mann DA, Lobel PS (1997) Propagation of damselfish (Pomacentridae) courtship sounds. J Acoust Soc Am 101:3783–3791 10.1121/1.418425

Mann DA, Lobel PS (1998) Acoustic behavior of the damselfish *Dascyllus albisella*: behavioral and geographic variation. Environ Biol Fish 51:421–428 10.1023/A:1007410429942

McCauley RD, Cato DH (2000) Patterns of fish calling in a nearshore environment in the Great Barrier Reef. Phil Trans R Soc B 355:1289–1293

McCormack L, Delikaris-Manias S, Farina A, Pinardi D, Pulkki V Real-time conversion of sensor array signals into spherical harmonic signals with applications to spatially localised sub-band sound-field analysis. In: 144th Convention of the Audio Engineering Society, Milan, Italy, 2018. pp 1–10

McElligott MM, Lammers MO (2021) Investigating spinner dolphin (*Stenella longirostris*) occurrence and acoustic activity in the Maui Nui Region. Front Mar Sci 8:703818 10.3389/fmars.2021.703818

McKenna MF et al. (2021) Advancing the interpretation of shallow water marine soundscapes. Front Mar Sci 8:719258 10.3389/fmars.2021.719258

McKenna MF et al. (2024) Understanding vessel noise across a network of marine protected areas. Environ Monit Assess 196:369 10.1007/s10661-024-12497-2

Minier L et al. (2025) Visualization and quantification of coral reef soundscapes using CoralSoundExplorer software. PLoS Comput Biol 21:e1012050 10.1371/journal.pcbi.1012050

Mooney TA et al. (2020) Listening forward: Approaching marine biodiversity assessments using acoustic methods. R Soc Open Sci 7:201287 10.1098/rsos.201287

Mouillot D, Graham NAJ, Villéger S, Mason NWH, Bellwood DR (2013) A functional approach reveals community responses to disturbances. Trends Ecol Evol 28:167–177 10.1016/j.tree.2012.10.004

Mouillot D et al. (2014) Functional over-redundancy and high functional vulnerability in global fish faunas on tropical reefs. Proc Natl Acad Sci USA 111:13757–13762 10.1073/pnas.1317625111

Mouy X, Black M, Cox K, Qualley J, Dosso S, Juanes F (2023) Identification of fish sounds in the wild using a set of portable audio-video arrays. Methods Ecol Evol 14:2165–2186 10.1111/2041-210X.14095

Muenzel D et al. (2024) Combining environmental DNA and visual surveys can inform conservation planning for coral reefs. Proc Natl Acad Sci USA 121:e2307214121 doi:10.1073/pnas.2307214121

Oliver SJ, Lobel PS (2013) Direct mate choice for simultaneous acoustic and visual courtship displays in the damselfish, Dascyllus albisella (Pomacentridae). Environ Biol Fish 96:447–457 10.1007/s10641-012-0028-z

Ozanich E, Thode A, Gerstoft P, Freeman LA, Freeman S (2021) Deep embedded clustering of coral reef bioacoustics. J Acoust Soc Am 149:2587–2601 10.1121/10.0004221

Parmentier E, Bertucci F, Bolgan M, Lecchini D (2021) How many fish could be vocal? An estimation from a coral reef (Moorea Island). Belg J Zool 151:1–29 10.26496/bjz.2021.82

Parmentier E, Lagardere JP, Vandewalle P, Fine ML (2005) Geographical variation in sound production in the anemonefish *Amphiprion akallopisos*. Proc R Soc B 272:1697–1703 10.1098/rspb.2005.3146

Parnell K, Merkens K, Huetz C, Charrier I, Robinson SJ, Pacini A, Bejder L (2024) Underwater soundscapes within critical habitats of the endangered Hawaiian monk seal: implications for conservation. Endang Species Res 54:311–329 10.3354/esr01336

Popper AN, Salmon M, Parvulescu A (1973) Sound localization by the Hawaiian squirrelfishes *Myripristis berndti* and *M. argyromus*. Anim Behav 21:86–97 10.1016/S0003-3472(73)80044-2

Pyć CD, Vallarta JH, Rice AN, Zeddies DG, Maxner EE, Denes SL (2021) Vocal behavior of the endangered splendid toadfish and potential masking by anthropogenic noise. Conserv Sci Pract 3:e352 10.1111/csp2.352

Raick X, Collet P, Under The Pole Consortium, Lecchini D, Bertucci F, Parmentier E (2023) Diel cycle of two recurrent fish sounds from mesophotic coral reefs. Sci Mar 87:e078 10.3989/scimar.05395.078

Rice AN, Farina SC, Makowski AJ, Kaatz IM, Lobel PS, Bemis WE, Bass AH (2022) Evolutionary patterns in sound production across fishes. Ichthyol Herpetol 110:1–12 10.1643/i2020172

Rountree RA, Gilmore RG, Goudey CA, Hawkins AD, Luczkovich JJ, Mann DA (2006) Listening to fish: Applications of passive acoustics to fisheries science. Fisheries 31:433– 446 10.1577/1548-8446(2006)31[433:LTF]2.0.CO;2

Ruiz NR (2026) Integrating video and acoustic non-invasive techniques to monitor coastal fish communities: A literature review with implications for MPA management. Acta Ethol 29:29 10.1007/s10211-026-00494-2

Salmon M (1967) Acoustical behavior of the menpachi, *Myripristis berndti*, in Hawaii. Pac Sci 21:364–381

Salmon M, Winn HE, Sorgente N (1968) Sound production and associated behavior in triggerfishes. Pac Sci 22:11–20

Sartori JD, Bright TJ (1973) Hydrophonic study of the feeding activities of certain Bahamian parrot fishes, Family Scaridae. Hydro Lab J 2:25–56

Schmidt RO (1986) Multiple emitter location and signal parameter estimation. IEEE Trans Antennas Propag 34:276–280 10.1109/tap.1986.1143830

Selkoe KA, Halpern BS, Toonen RJ (2008) Evaluating anthropogenic threats to the Northwestern Hawaiian Islands. Aquat Conserv Mar Freshw Ecosyst 18:1149–1165 10.1002/aqc.961

Smith MK (1993) An ecological perspective on inshore fisheries in the main Hawaiian Islands. Mar Fish Rev 55:34–49

Stevenson RA (1963) Life history and behavior of Dascyllus albisella (Gill), a pomacentrid reef fish. Ph.D. Dissertation, University of Hawai’i at Manoa

Sugai LSM, Balantic C, Clink DJ, Ramesh V, Kahl S, Klinck H, Wood CM (2026) Acoustic indices are not useful for biodiversity research. Methods Ecol Evol 17:1506–1518 10.1111/2041-210x.70285

Thode AM et al. (2021) Automated two-dimensional localization of underwater acoustic transient impulses using vector sensor image processing (vector sensor localization). J Acoust Soc Am 149:770–787 10.1121/10.0003382

Tricas TC, Boyle KS (2014) Acoustic behaviors in Hawaiian coral reef fish communities. Mar Ecol Prog Ser 511:1–16 10.3354/meps10930

Tricas TC, Boyle KS (2021) Parrotfish soundscapes: implications for coral reef management. Mar Ecol Prog Ser 666:149–169 10.3354/meps13679

van Oosterom L, Montgomery JC, Jeffs AG, Radford CA (2016) Evidence for contact calls in fish: conspecific vocalisations and ambient soundscape influence group cohesion in a nocturnal species. Sci Rep 6:19098 10.1038/srep19098

Williams B et al. (2022) Enhancing automated analysis of marine soundscapes using ecoacoustic indices and machine learning. Ecol Indic 140:108986 10.1016/j.ecolind.2022.108986

Williams B et al. (2025) Using tropical reef, bird and unrelated sounds for superior transfer learning in marine bioacoustics. Phil Trans R Soc B 380:20240280 10.1098/rstb.2024.0280

Wolfe K et al. (2020) Priority species to support the functional integrity of coral reefs Oceanogr Mar Biol Ann Rev 58:179–318 10.1201/9780429351495-5

Zurk LM, Ou HH, Schecklman S, Lutwak A (2014) Acoustic monitoring of marine conservation areas. Mar Technol Soc J 48:21–32 10.4031/MTSJ.48.6.7

